# Menin inhibition impairs metastatic colonization of Ewing sarcoma

**DOI:** 10.1101/2025.11.10.687648

**Authors:** Katherine A. Braun, Nicolas M. Garcia, Mohamed Ahmed, Darleen S. Tu, Stephanie I. Walter, Emma D. Wrenn, Megan E.B. Dean, Elizabeth R. Lawlor

## Abstract

Menin is a scaffolding protein that interacts with context-specific partners to regulate gene expression. In MLL-rearranged leukemias, Menin:MLL interactions drive leukemogenesis and Menin inhibitors have been FDA approved for these cancers. We previously reported that Menin promotes oncogenic phenotypes in Ewing sarcoma (EwS). Here, we sought to define EwS-specific functions of Menin and determine if Menin inhibitors could be therapeutically leveraged for these tumors. Genetic knockout of Menin had no impact on EwS cell proliferation in vitro but metastatic potential of Menin-depleted cells in vivo was impaired. Transcriptional profiling of Menin knockout cells in vitro showed reproducible downregulation of MYC signature genes and upregulation of developmental programs. Conversely, transcriptional rewiring of developmental genes and restoration of MYC target gene expression were evident in tumors that arose from Menin knockout cells. Exposing EwS cells to the Menin inhibitor VTP50469 (revumenib) inhibited expression of MYC targets and co-immunoprecipitation studies detected Menin:MYC interactions that were partially disrupted by the drug. Metastatic colonization of disseminated EwS cells in vivo was significantly inhibited in mice fed VTP50469 chow. Together these findings implicate Menin as a mediator of EwS metastasis and suggest that Menin inhibitors warrant investigation as novel therapeutics for patients with high-risk disease.

## Introduction

The *MEN1* (Multiple Endocrine Neoplasia 1) gene encodes Menin, a unique scaffolding protein that regulates many cellular processes including transcription, DNA damage, cell cycle and signal transduction through interactions with a wide range of context-dependent partner proteins (1, 2). Menin has broad and fundamental roles in early development. *Men1* null mice are early embryonic lethal at E11.5-E13.5 due to widespread defects in organogenesis (3). In addition, studies of human embryonic stem cells implicate Menin in regulating timing of endoderm, cardiomyocyte and neuronal differentiation (4). Targeted deletion of Menin from embryonic murine neural crest cells and somite precursors disrupts cranio-facial skeleton and rib development in vivo (5) and misregulation of Menin expression in mesenchymal progenitor cells in vitro disrupts lineage commitment and terminal differentiation (6–8). Thus, Menin plays diverse roles in regulating fidelity of organogenesis and tissue differentiation during normal development. Importantly, the diverse and highly tissue-specific functions of Menin are determined by its context-dependent interaction partners, including epigenetic modifiers (e.g. MLL1, MLL2), transcription factors (RUNX2, JUND), and signaling pathway components (1, 6, 8–11).

In human disease, *MEN1* was first identified as a tumor suppressor gene: germline inactivation of *MEN1* is associated with the genetic cancer predisposition syndrome multiple endocrine neoplasia (12–14). Patients with MEN1 syndrome are predisposed to developing parathyroid and pituitary gland tumors, neuroendocrine tumors of the pancreas, as well as other endocrine and non-endocrine (e.g. meningioma) cancers (15). By contrast in mixed-lineage leukemia (MLL)-rearranged (MLLr) leukemia, Menin functions as an oncogene (10, 16). Protein:protein interactions between Menin and MLL fusion proteins drive transcriptional activation of leukemogenic genes. Significantly, pharmacologic inhibitors of Menin have been developed that disrupt Menin interactions with MLL1-fusions and with wild-type MLL1 (*KMT2A*) and these agents induce terminal differentiation and subsequent death of MLLr and NPM1-mutant (NPM1m) leukemia (17–19). Revumenib (Syndax Pharmaceuticals, Inc., USA) was recently granted FDA approval for treatment of patients with MLLr and NPM1m leukemia (20) and multiple additional Menin inhibitors are currently in development (21). Significantly, recent studies have also identified pro-tumorigenic roles for Menin, and/or Menin:MLL in some solid tumors. These include carcinomas such as breast, lung, endometrial, ovarian, colorectal, and prostate cancer and tumors of mesenchymal origin such as gastrointestinal stromal tumors and Ewing sarcoma (EwS) (Reviewed in (22)). If and how Menin inhibitors could be leveraged for solid tumor therapies requires that tumor type-specific functions of Menin and its interaction partners be elucidated.

EwS tumors are aggressive cancers of bone and soft tissues that are of presumed mesenchymal and/or neural crest stem cell origin (23). Multiagent systemic chemotherapy combined with local control can induce long term remission in most patients who present with localized disease (23, 24). However, in up to 25% of cases, tumors recur at distant sites and patients who relapse with metastatic disease are rarely cured. We previously reported that Menin protein is highly expressed by EwS cells and that partial inhibition of Menin blunts tumorigenicity and pro-oncogenic metabolic programs (25–27). For the current study, we generated Menin knockout EwS cells to definitively characterize how Menin contributes to EwS pathogenesis. Our results show that Menin is required for colonization and metastatic competence of EwS cells in vivo. Mechanistically, transcriptomic profiling revealed reproducible downregulation of nucleic acid biosynthesis programs and MYC target genes in *MEN1*-knockout (MEN1-KO) cells. Conversely, Menin depletion led to upregulated expression of mesenchymal differentiation and development genes. Significantly, tumors that successfully arose from MEN1-KO cells were transcriptionally rewired to reverse these effects: expression of MYC target genes was restored while TGFβ and epithelial mesenchymal transition (EMT) gene programs were suppressed. Thus, these data suggest that high levels of Menin in EwS cells promote tumorigenicity and metastatic colonization, at least in part, by promoting transcriptional activity of MYC. It was previously reported that in HT-1080 fibrosarcoma cells, Menin directly interacts with MYC at E-boxes to amplify transcription of MYC-bound target genes (28). We performed co-immunoprecipitation studies and confirmed that Menin and MYC interact in EwS cells. Moreover, our data show that this interaction occurs outside Menin/MLL-containing compass-like complexes. We next exposed EwS cells to VTP50469, the preclinical precursor to revumenib, to test the impact of Menin:MLL interaction inhibitors on Menin:MYC interactions and MYC target gene expression. Notably, VTP50469 partially disrupted the interaction between Menin and MYC and significantly blunted the MYC signature. Given the profound impact of Menin loss on metastatic colonization, we tested if VTP50469 would inhibit outgrowth of disseminated EwS tumor cells into macroscopic tumors. Feeding mice therapeutic doses of VTP50469 chow significantly reproducibly inhibited colonization of disseminated tumor cells.

Together these data confirm the critical importance of Menin to tumorigenic phenotypes in EwS and suggest that Menin inhibitors could be repurposed to inhibit tumor colonization in patients who are at high risk of metastatic relapse.

## Results

### In vivo tumorigenicity of MEN1-KO EwS cells is impaired

To examine the function of Menin in EwS, we generated Menin knockout (MEN1-KO) cells using the CRISPR/Cas9 system. Two guide RNAs, one targeting exon 2 (guide 1) and one targeting exon 3 (guide 5) were generated and used to knock out the *MEN1* gene in two EwS cell lines, A673 and TC32 (**Figure 1A**). In parallel, control cell lines were created using a non-targeting gRNA. Single cell clones were isolated and propagated to generate clonally-derived MEN1-KO and control cell lines. *MEN1* transcript levels were reduced in all MEN1-KO clones (**Figure 1B**) and Menin protein was not detected by western blot, confirming complete loss of the protein (**Figure 1C**). Interestingly, protein levels of both MLL1 (*KMT2A*) and MLL2 (*KMT2B*) were reproducibly reduced in MEN1-KO cells indicating that both MLL1 and MLL2 proteins are highly expressed by EwS cells and expression is maintained by Menin.

**Figure 1.**
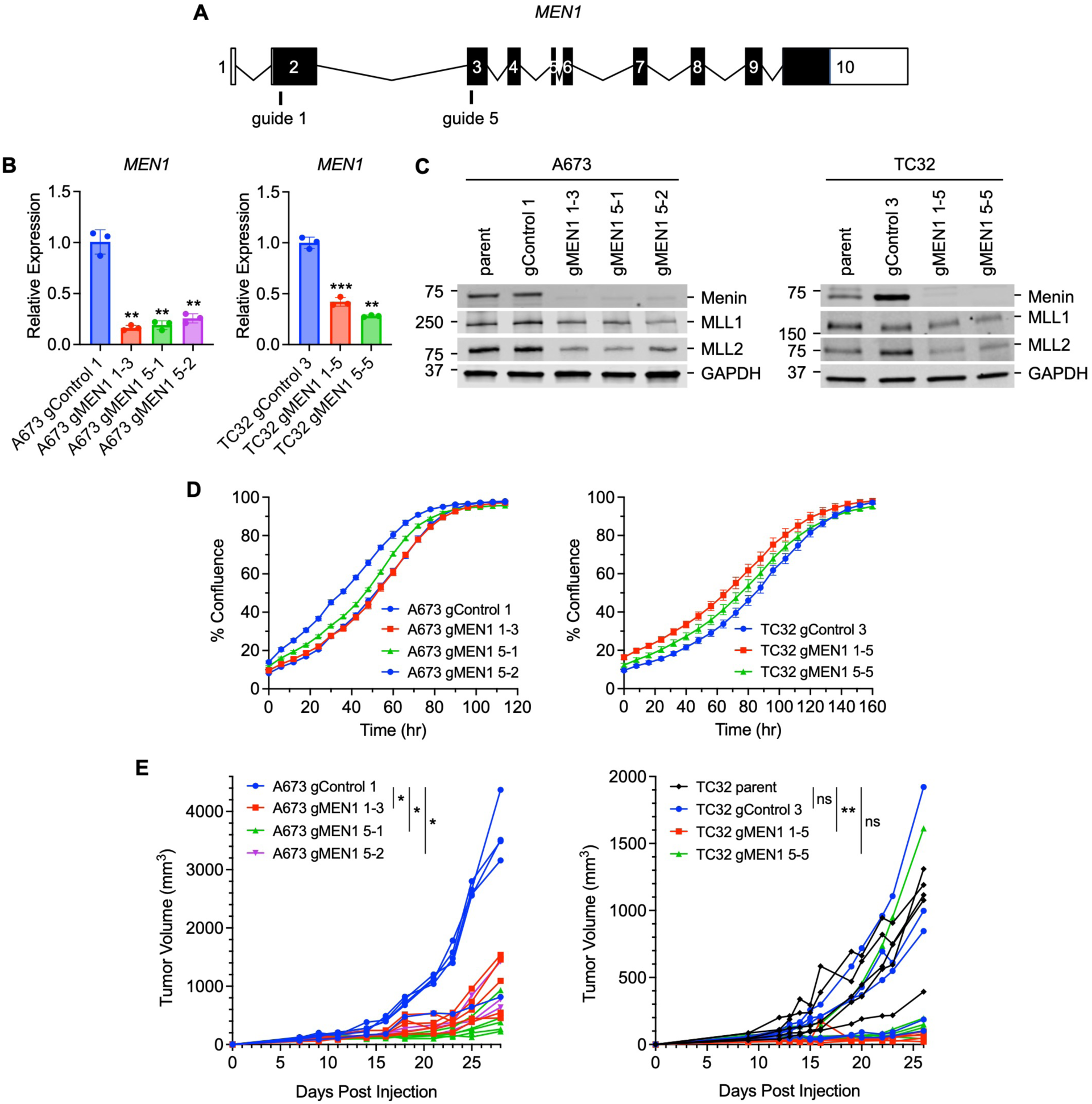
Menin promotes EwS tumorigenesis. (**A**) CRISPR guides in exons 2 (guide 1) and 3 (guide 5) used to knock out *MEN1*. (**B**) *MEN1* expression in control and MEN1-KO clones. RT-qPCR determined expression levels and p-values plotted relative to control cell lines. (**C)** Western blot of Menin, MLL1, MLL2 and GAPDH in control and MEN1-KO clones. (**D**) Incucyte proliferation assays of control and MEN1-KO clones plotting percent confluence over time. Representative of n=3-4. (**E**) Subcutaneous tumor growth of A673 and TC32 control and MEN1-KO cells in NSG mice. Tumor volume is plotted over time for each mouse. The mean tumor size in mice from each cell line at endpoint was used to determine p values. ns p > 0.05, * p ≤ 0.05, ** p ≤ 0.01, *** p ≤ 0.001.

To assess whether loss of Menin affected growth of EwS cells in vitro we used Incucyte assays to measure real-time proliferation. As shown, proliferation of MEN1-KO cells was not significantly changed relative to controls (**Figure 1D**). Next, we evaluated the in vivo tumorigenic potential of MEN1-KO cells. Notably, despite an absence of observable effects on cell proliferation in vitro, the capacity of A673 and TC32 MEN1-KO cells to generate subcutaneous tumors in NOD scid gamma (NSG) mice was significantly impaired compared to both parental and clonally-derived control gRNA cell lines (**Figure 1E**). Thus, depleting EwS cells of Menin inhibits their ability to initiate local tumor formation, confirming that Menin plays a critical role in mediating tumorigenicity.

### Loss of Menin leads to widespread transcriptional changes in cancer-associated gene programs

Menin has well-established roles in regulating gene transcription in development and in cancer. Therefore, to identify Menin-regulated genes in EwS, we performed transcriptomic profiling of control and MEN1-KO A673 and TC32 cells. RNA-seq data showed widespread and highly significant changes in gene expression in both A673 and TC32 MEN1-KO cell lines compared to controls, revealing a critical role for Menin as a master regulator of gene transcription (**Supplemental Figure 1**). To identify changes that were most likely to mediate loss of tumorigenicity, we focused on genes that were reproducibly altered upon Menin loss in both cell lines. Despite evidence of cell context-specificity, 1124 transcripts were commonly downregulated (**Figure 2A**) and 1258 genes were upregulated (**Figure 2B**) in both models. Given the breadth of shared genes, we applied Hallmark and Gene Ontology Biological Processes (GOBP) gene set enrichment tools to the gene lists to identify key biologic programs that were reproducibly altered in MEN1-KO cells. mTORC1 signaling, E2F targets, MYC targets, and G2-M checkpoint genes were highly enriched among genes that were downregulated in MEN1-KO cells (i.e. Menin-activated genes) (**Figure 2C**). GOBP analysis of Menin-activated genes further labeled enrichment of nucleotide metabolism, a key process in cell growth and proliferation. Though no changes in proliferation were detected in vitro, growth of MEN1-KO cells was inhibited in vivo (**Figure 1**), nominating the tumor microenvironment (TME) as a player in mediating the effects of Menin on cell proliferation. Genes reproducibly upregulated in MEN1-KO cells (i.e. Menin-repressed genes) were enriched for mesenchymal developmental programs including EMT and myogenesis, as well as the apical junction pathway that regulates cell-cell contacts (**Figure 2D**). Biological processes involved in cell migration and modulation of extracellular matrix (ECM) and the tumor microenvironment (TME) were enriched in Menin-repressed genes. Consistent with this, MEN1-KO cells were more invasive than controls when embedded in 3D collagen (Figure 2E and **F**). Thus, these findings suggest that Menin maintains the tumorigenicity of EwS cells by sustaining activation of growth-promoting programs and by suppressing differentiation programs. Moreover, the impact of these Menin-regulated genes on tumor behavior is minimal in standard 2D culture but highly evident in the context of an in vivo TME.

**Figure 2.**
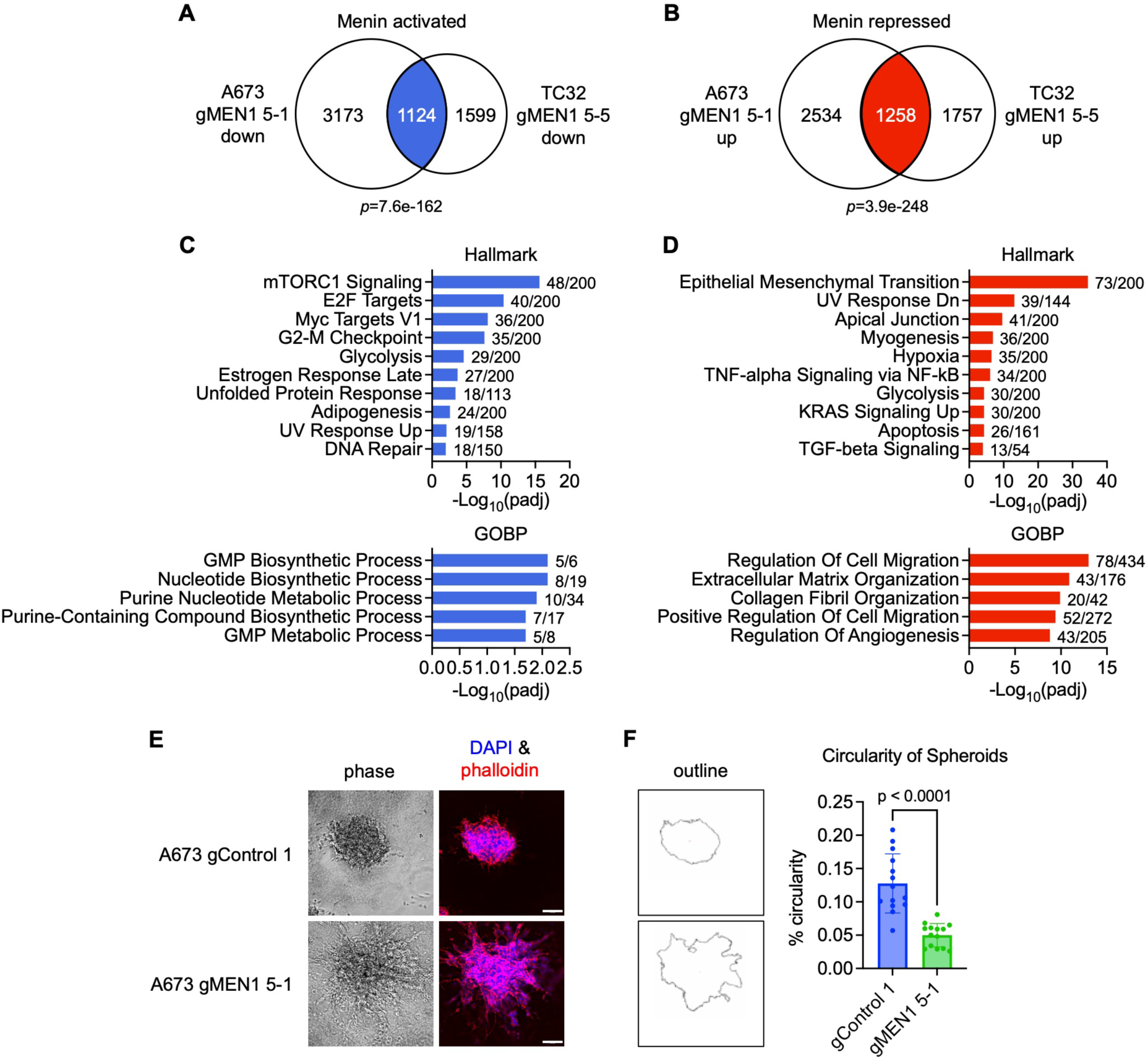
MEN1-KO EwS cells have altered expression of proliferation and differentiation programs. (**A** and **B**) Overlap of downregulated (**A**) and upregulated (**B**) genes in A673 and TC32 MEN1-KO cells. Significant genes in each model were defined as padj<0.05 as described in methods. (**C** and **D**) The top 10 enriched Hallmark pathways and the top 5 enriched GOBP pathways among overlapping downregulated (**C**) and upregulated (**D**) genes as determined by Enrichr. The number of overlapping/total genes is shown for each pathway. (**E**) Invasion of control and MEN1-KO cells embedded in rat tail collagen for 5 days. Representative phase and phalloidin (red)/DAPI (blue) stained images are shown. (**F**) Invasion in collagen was quantified and compared between control and MEN1-KO cells using spheroid circularity, as described in methods (scale bars=100 µm).

### Menin is required for metastatic competence of EwS cells

EwS cells are highly heterogeneous and exist along a transcriptional spectrum from highly undifferentiated to more mesenchymal cell states (29–31). EwS cells that upregulate mesenchymal genes show enhanced capacity to metastasize in vivo. Given our discovery that EMT genes and invasion were augmented in MEN1-KO cells in vitro, we next tested their metastatic competence in vivo. We first evaluated metastatic colonization using tail vein models. Despite their more mesenchymal and invasive profiles, MEN1-KO cells were largely incapable of generating macroscopic tumors when intravenously administered to recipient mice. This was evident by serial luciferase imaging (**Figure 3A**) and at necropsy (**Figure 3B**). Whereas control cells generated numerous tumors in the livers of NSG mice that nearly completely effaced normal liver, MEN1-KO cells generated few to no visible tumors (**Figure 3B**).

**Figure 3.**
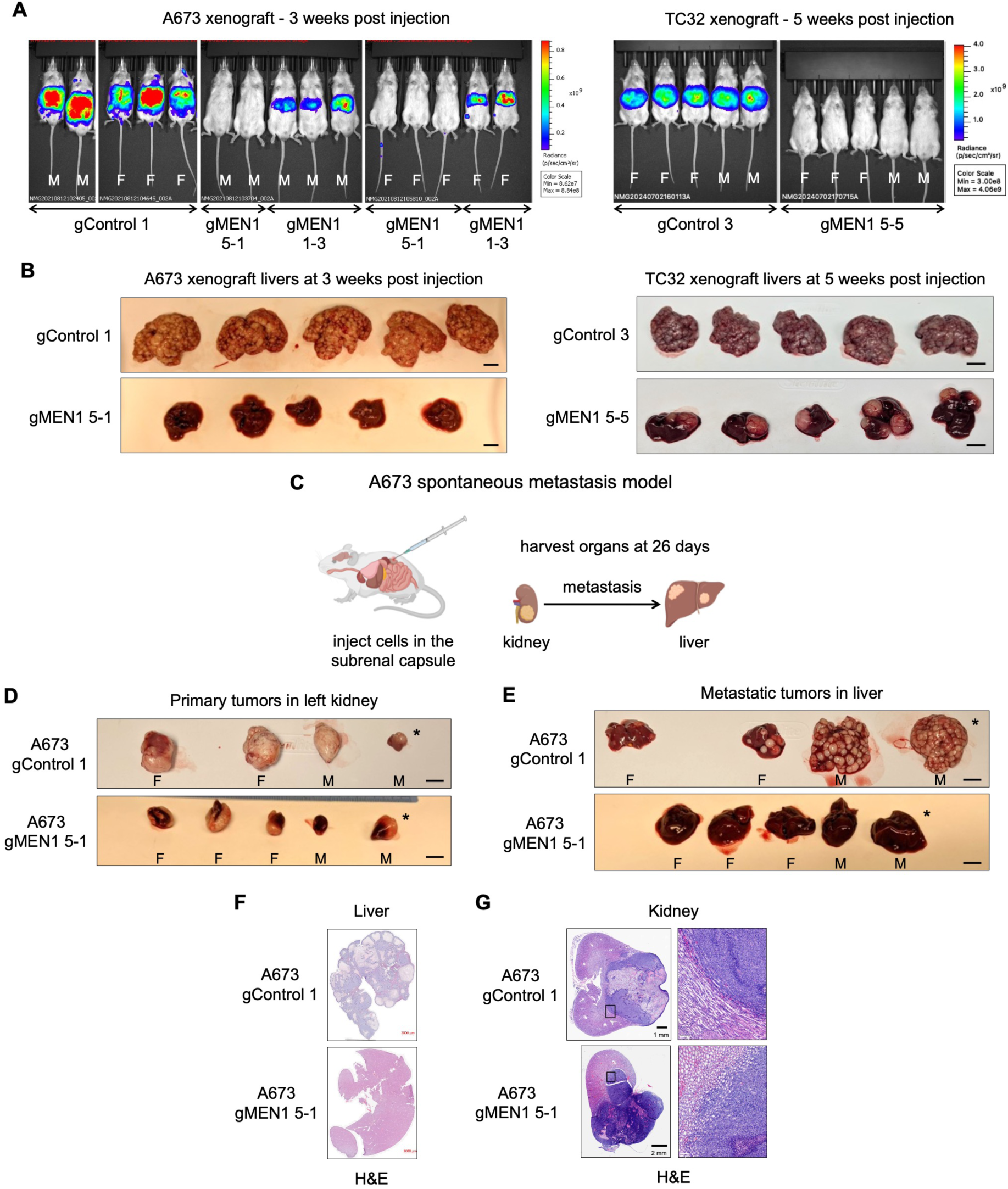
Menin promotes colonization and metastasis. (**A**) IVIS images of NSG mice 3- or 5-weeks post-tail vein inoculation with 1e6 control or MEN1-KO luciferase-tagged A673 and TC32 cells, respectively. Sex of mice are indicated as M for male and F for female. (**B**) Photos of livers at necropsy show markedly diminished macroscopic tumor burden in mice that received MEN1-KO vs control cells. (**C)** Diagram of the spontaneous metastasis model. 2e5 A673 control or MEN1-KO cells were injected in the subrenal capsule of the left kidney of NSG mice and 26 days post-injection the left kidney and the liver were harvested. (**D** and **E**) Photographic images of left kidney (**D**) and liver (**E**) from each mouse are shown (scale bars=1 cm). H&E stained sections of the livers (**F**) and kidneys (**G**) from organs marked by asterisks (*) in **D** and **E**. Right panel in (**G**) is 10x amplified image to show tumor-kidney border.

Having established that colonization of circulating MEN1-KO cells was inhibited in the tail vein model, we next used a subrenal capsule model to study spontaneous metastasis of EwS cells from kidney to liver. A673 cells were injected by ultrasound guidance into the subrenal capsule of the left kidney, as we have previously described (**Figure 3C**) (32). Control (N=4) and MEN1-KO (N=5) cell recipient mice were euthanized after 26 days, at which point all mice had developed kidney tumors. In keeping with subcutaneous tumor models, tumors derived from MEN1-KO cells were mainly smaller than control tumors (**Figure 3D**). Significantly, macroscopic tumors were detected in the livers of all control and none of the MEN1-KO cell recipient mice (**Figure 3E**). Importantly, this defect in spontaneous metastasis of MEN1-KO cells could not be entirely explained by the smaller size of the primary tumors. As shown (tumors marked by * in Figure 3D and **E**), a control tumor of comparable size to MEN1-KO tumors generated dozens of metastases that completely replaced normal liver. The marked difference in liver tumor burden, in conjunction with comparable primary tumor size, were confirmed by H&E staining (**Figure 3F**). Interestingly, despite their evident loss of metastatic competence, MEN1-KO cells successfully invaded normal adjacent kidney parenchyma (**Figure 3G**). Thus, these in vivo studies of MEN1-KO cells collectively demonstrate that metastatic competence of EwS cells is profoundly dependent on Menin. In particular, they suggest that Menin is required for successful colonization of EwS cells at metastatic sites.

### Tumors derived from MEN1-KO cells are transcriptionally rewired to restore Menin-regulated gene programs

Despite a marked delay in engraftment, some A673 MEN1-KO cells did eventually generate tumors in vivo despite persistent loss of Menin protein (**Figure 1E**, **Figure 4A, and Supplemental Figure 2**). We hypothesized that defining the mechanisms by which these MEN1-KO cells had bypassed the effects of Menin loss would illuminate the most critical transcriptional targets of Menin that mediate its effects on tumor colonization. To address this, we performed RNA-seq on 12 tumors: three generated from the control gRNA cell line and three from each of the three MEN1-KO cell lines (marked by * in **Figure 4A**). Comparison of tumor transcriptomes revealed thousands of differentially expressed transcripts that were up- or downregulated in MEN1-KO tumors compared to controls (**Figure 4B and C**). Focusing on the transcripts that were reproducibly altered in tumors from all three MEN1-KO cell lines identified 936 commonly upregulated and 1059 downregulated genes. Commonly upregulated genes were enriched for G2-M checkpoint, E2F and MYC targets (**Figure 4D**). Remarkedly, these same pathways were significantly downregulated in MEN1-KO cells (**Figure 2C**). Comparing expression patterns of specific genes in these over-represented pathways confirms that expression of core oncogenic genes was reestablished in MEN1-KO cells coincident with tumor outgrowth (**Figure 4E**). Similarly, transcriptional rewiring of Menin-repressed genes was also evident in MEN1-KO cell-derived tumors (**Figure 4F**).

**Figure 4.**
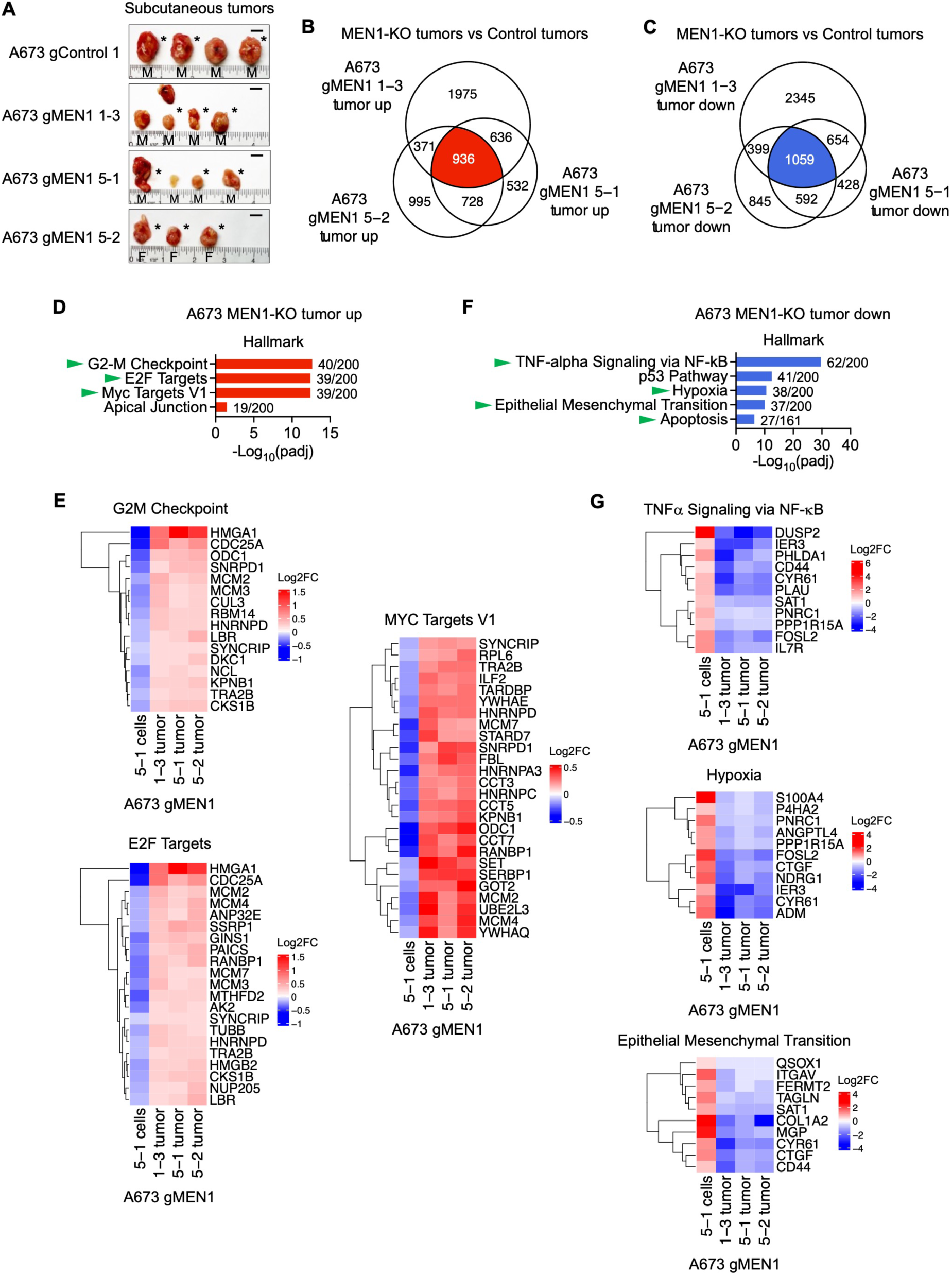
Tumors derived from MEN1-KO cells are transcriptionally rewired. (**A**) Photos of tumors removed from NSG mice 28 days after subcutaneous injection of A673 control and MEN1-KO cells (from Figure 1E). Three tumors from each cell line were processed for RNA-seq (denoted by *). Sex of mice is indicated (M for male, F for female). Scale bars=1cm. (**B** and **C**) Overlap of genes that were significantly upregulated (**B**) or downregulated (**C**) (padj<0.05) in MEN1-KO cell-derived tumors compared to controls. (**D**) Enriched Hallmark pathways (padj<0.05) as determined by Enrichr among shared upregulated genes in MEN1-KO tumors. Overlap/total genes shown for each pathway. Green arrows indicate pathways that were identified as being downregulated in A673 MEN1-KO cells in vitro (see Figure 2C). (**E**) Heatmap of log2FC of genes from Hallmark pathways that were downregulated in MEN1-KO cells in vitro and upregulated in MEN1-KO cell-derived tumors in vivo. (**F**) Enriched Hallmark pathways (padj<0.05) as determined by Enrichr among shared downregulated genes in MEN1-KO tumors. Overlap/total genes shown for each pathway. Green arrows indicate pathways that were identified as being upregulated in A673 MEN1-KO cells in vitro (see Figure 2D). (**G**) Heatmap of log2FC of genes from Hallmark pathways that were upregulated in MEN1-KO cells in vitro and downregulated in MEN1-KO cell-derived tumors in vivo.

Restoration of tumorigenicity was associated with renewed suppression of genes that were relatively upregulated in MEN1-KO cells (**Figure 2D**), most predominantly genes that regulate NF-κB, hypoxia, and EMT pathways (**Figure 4F and G**). Thus, these studies of in vivo tumors confirm that Menin serves as a master regulator of oncogenic gene programs in EwS. Moreover, they reinforce the hypothesis that Menin-dependent activation of MYC target gene expression is critical for maintenance of tumorigenicity.

### Menin and MYC proteins interact outside MLL-complexes in EwS cells

Our transcriptomic analyses identified MYC target genes as being among the most highly and reproducibly altered by Menin depletion. We therefore sought to address whether this is mediated via direct regulation of MYC levels. Quantitative RT-PCR and RNA-seq analyses revealed no change in *MYC* mRNA expression in either MEN1-KO cells or tumors (**Figure 5A and B**). Likewise, MYC protein levels were largely unchanged, although expression was modestly increased in a few tumors derived from MEN1-KO cells (**Figure 5C and D**). Thus, these data suggest that Menin does not activate the MYC signature by directly inducing expression of MYC itself. Importantly, it was previously reported that Menin can directly interact with MYC and that protein:protein interactions between Menin and MYC at E-boxes leads to RNA polymerase II and P-TEFb recruitment and transcriptional amplification (28). To assess whether Menin and MYC interact in EwS cells, we performed co-immunoprecipitation experiments (co-IP). As shown (**Figure 5E** and Supplemental Figure 3), MYC was pulled down by the Menin antibody in A673, CHLA10, and TC32 EwS cells but not in U2OS osteosarcoma cells. By contrast, the H3K4 methyltransferase complex proteins MLL2 and WDR5 co-immunoprecipitated with Menin in all cell lines. These findings confirm that, unlike its canonical trithorax complex partners, Menin interactions with MYC are context specific. Next, we reversed the IP antibodies to validate that MYC can also pull down Menin. These studies confirmed interactions between MYC and Menin, along with its canonical binding partner MAX (**Figure 5F**). Conversely, MYC did not pull down either MLL2 or WDR5, revealing that Menin interactions with MYC occur outside of MLL-containing methyltransferase complexes. These data suggest that Menin-dependent activation of the MYC signature in EwS is mediated by Menin:MYC interactions. Whether this is orchestrated by selective recruitment of RNA polymerase II to MYC bound E-boxes, or other EwS-specific mechanisms, requires additional investigation.

**Figure 5.**
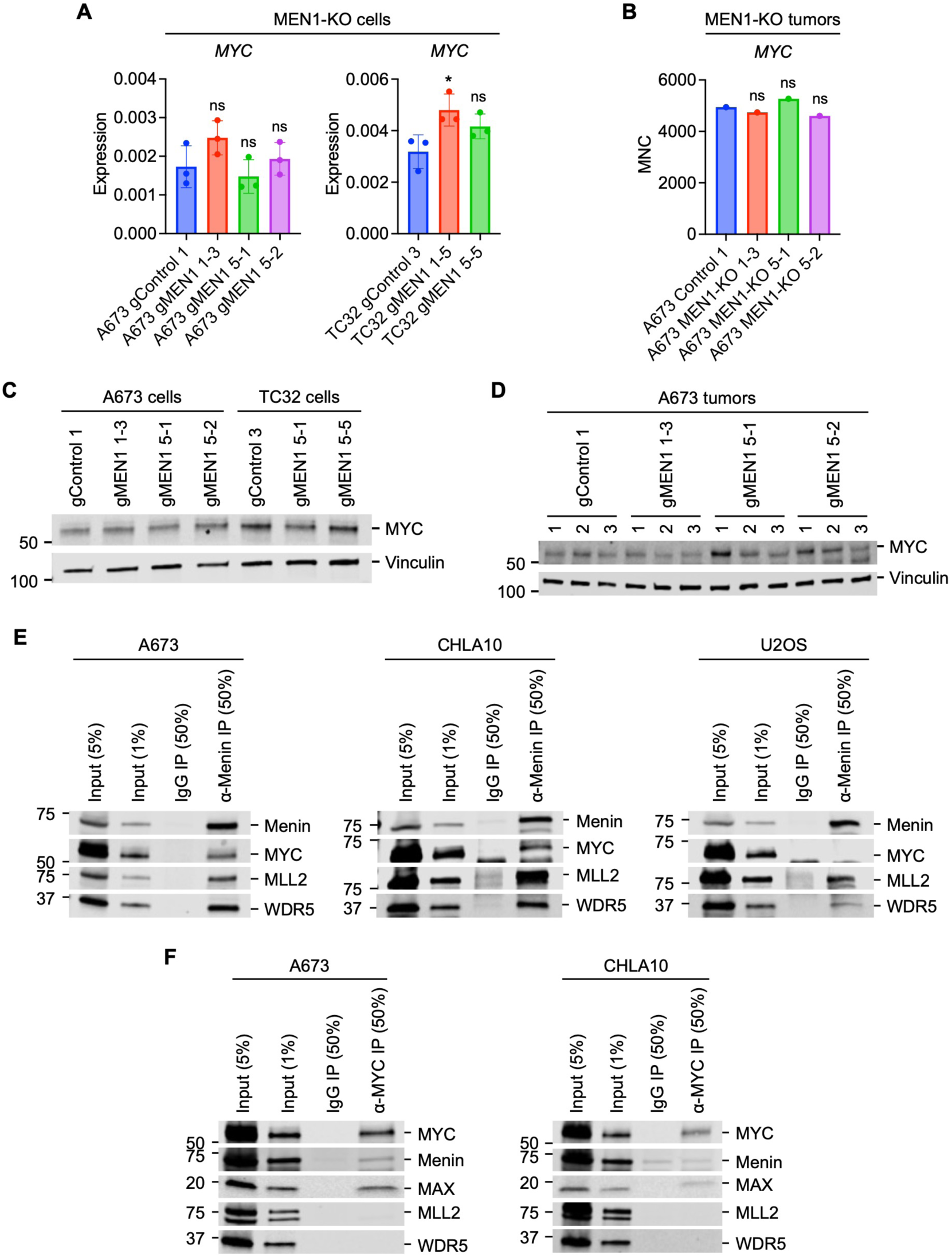
Menin and MYC form a complex in EwS cells. (**A**) *MYC* transcript levels in control and MEN1-KO cells as determined by RT-qPCR. ns p > 0.05, * p ≤ 0.05. (**B**) *MYC* transcript counts from RNA-seq of A673 control and MEN1-KO tumors. (**C**) Western blot of MYC and Vinculin levels in A673 and TC32 control and MEN1-KO cells (**D**) Western blot of MYC and Vinculin in tumors derived from A673 control and MEN1-KO cells. (**E**) Co-immunoprecipitations with an anti-Menin antibody were performed on A673, CHLA10 and U2OS nuclear extracts and immunoblotted for Menin, MYC, MLL2 and WDR5. (**F**) Co-immunoprecipitations with an anti-MYC antibody were performed on A673 and CHLA10 nuclear extracts and immunoblotted for MYC, Menin, MAX, MLL2 and WDR5.

### Pharmacologic inhibition of Menin downregulates expression of MYC targets

Studies of MEN1-KO cells identified Menin as a master regulator of MYC targets and other pro-oncogenic gene programs in EwS. We next sought to test whether pharmacologic inhibitors of Menin could in whole or in part reverse this activity. For these investigations we utilized VTP50469, the pre-clinical precursor of revumenib which was recently FDA approved for treatment of MLLr and NPM1m leukemia. Importantly, exposing EwS cells to VTP50469 had no effect on Menin transcript (**Supplemental Figure 4A**) or protein expression (**Supplemental Figure 4B**) allowing us to compare the impact of pharmacologic loss of function to the effects of protein loss. Consistent with genetic knockout, pharmacologic inhibition of Menin had no impact on EwS cell proliferation in vitro (**Figure 6A**). However, unlike MEN1-KO cells, VTP50469-treated cells did not exhibit augmented invasion in 3D collagen (**Figure 6B and C**) indicating that Menin-inhibition using a Menin:MLL interaction inhibitor does not completely phenocopy the effects of Menin depletion.

**Figure 6.**
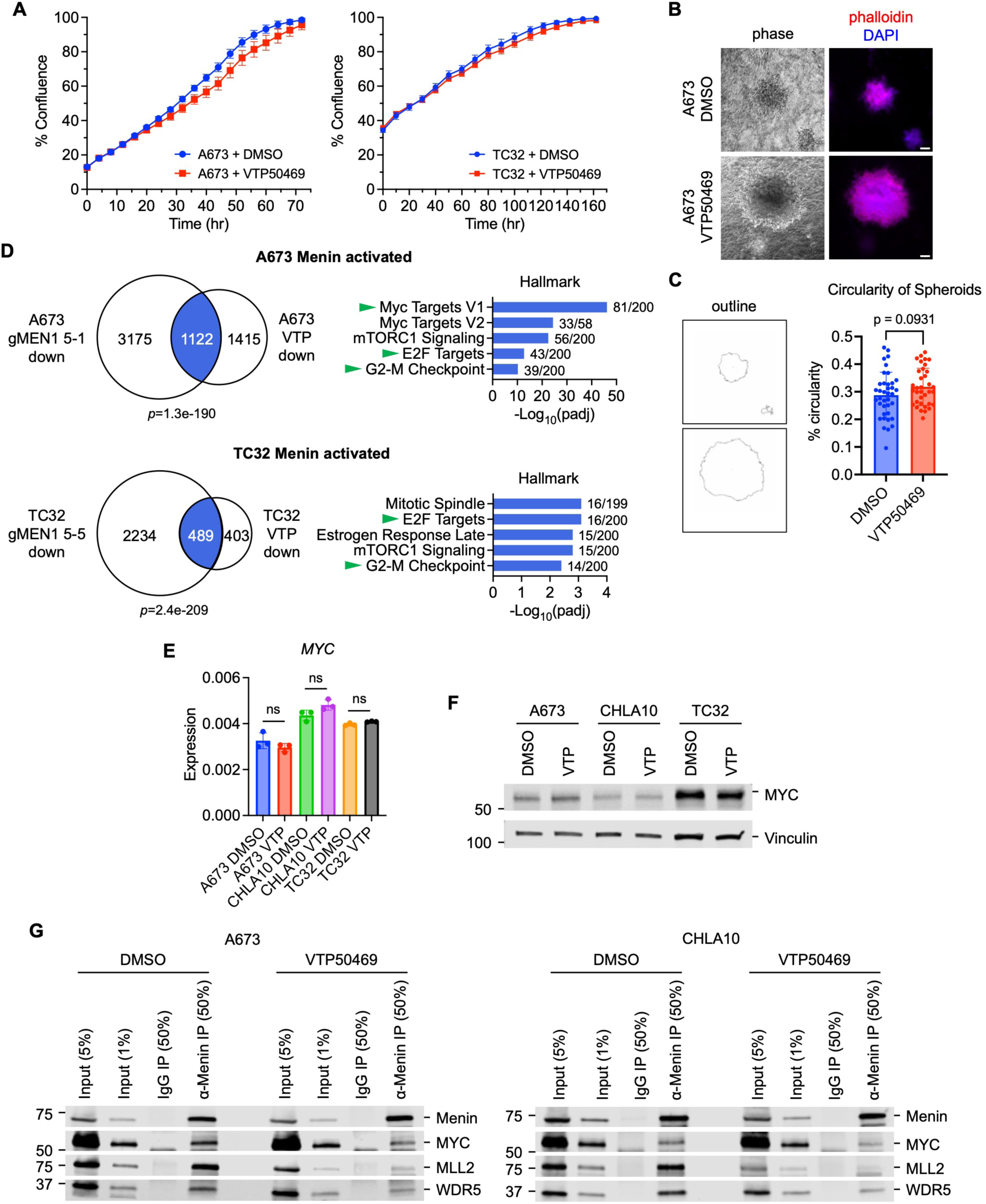
Menin inhibition regulates key tumorigenic pathways. (**A**) Incucyte proliferation assay plotting percent confluence for A673 and TC32 cells treated with 0.1% DMSO or 10 µM VTP50469 starting at time 0. Representative of n=3-4). (**B** and **C**) Invasion of cells from a sphere of A673 cells embedded in rat tail collagen and treated with 0.1% DMSO or 10 µM VTP50469 for 5 days. (**B**) Representative phase and phalloidin (red)/DAPI (blue) stained images are shown (scale bars=100 µm). (**C**) Plot of circularity quantified from replicate spheroids treated as in (**B**). (**D**) A673 and TC32 cells were treated with 0.1% DMSO or 10 µM VTP50469 for 72 hours and genes that were significantly downregulated in VTP50469-treated cells (padj<0.05) were overlapped with genes downregulated in MEN1-KO cells. The top 5 enriched Hallmark pathways for the overlapping downregulated genes in each cell line are shown along with the overlap/total genes for each pathway. The green arrows indicate pathways that were reactivated in A673 MEN1-KO tumors (see Figure 4D). (**E**) *MYC* transcript levels in A673, CHLA10 and TC32 cells treated with 0.1% DMSO or 10 µM VTP50469 for 72 hours as determined by RT-qPCR (ns p > 0.05). (**F**) Western of MYC and Vinculin in cells treated with 0.1% DMSO or 10 µM VTP50469 for 72 hours. (**G**) Co-immunoprecipitations with an anti-Menin antibody were performed on nuclear extracts from A673 and CHLA10 cells treated with 0.1% DMSO or 10 µM VTP50469 for 72 hours and immunoblotted for Menin, MYC, MLL2 and WDR5.

We next evaluated the effects of VTP50469 on gene transcription. Despite having no obvious effects on in vitro cell phenotypes, Menin inhibition led to up- or downregulation of thousands of genes (**Supplemental Figure 4C and D**). Though highly significant differences were observed between two different EwS cell lines, genes involved in cell signaling and cell cycle regulation were commonly over-represented among downregulated genes (**Supplemental Figure 4E**) while metabolic and developmental regulators were enriched in upregulated genes (**Supplemental Figure 4F**). To identify overlapping transcriptional effects of genetic and pharmacologic Menin inhibition strategies we compared VTP50469-treated to MEN1-KO cells. Intriguingly, like Menin depletion, VTP50469 significantly suppressed expression of MYC target genes (**Figure 6D**). Moreover, this effect of VTP50469 occurred without any significant changes to MYC transcript or protein (**Figure 6E and F**). Thus, we used co-IP studies to test if the drug inhibited interactions between Menin and MYC. As expected, and consistent with its mechanism of action as a Menin:MLL interaction inhibitor, VTP50469 dramatically and reproducibly reduced Menin-dependent pull down of both MLL2 and WDR5 proteins (**Figure 6G**). Significantly, VTP50469-treated cells also showed diminished interactions between Menin and MYC (**Figure 6G**).

Thus, pharmacologic disruption of Menin:MLL interactions in EwS cells leads to downregulated expression of pro-oncogenic MYC target genes and this is associated with partial disruption of protein:protein interactions between Menin and MYC.

### Menin inhibition reduces metastatic colonization in vivo

Our studies of MEN1-KO cells demonstrated that Menin is required for successful metastatic colonization of EwS tumor cells in vivo. In addition, transcriptomic profiling of MEN1-KO cells and tumors identified amplification of MYC targets as a likely promoter of Menin-dependent oncogenic states. Given our findings that VTP50469 disrupts Menin:MYC interactions and blunts expression of MYC target genes in vitro, we sought to test if the drug could be leveraged to block tumor colonization in vivo. To achieve this, we designed a model of disseminated micrometastatic disease that would allow comparison of metastatic outgrowth over time in control and VTP50469-treated NSG mice (**Figure 7A**). GFP-luciferase-labeled A673 cells were injected by tail vein and IVIS imaging one-hour post-cell injection confirmed that tumor cells had successfully entered the systemic circulation. One week post cell injections, no tumor was detectable by IVIS in any mice and animals were randomized to control or treatment groups. Inhibitor-treated mice were fed 0.1% VTP50469 chow (Syndax Pharmaceuticals, Inc., USA), a dose that effectively inhibits Menin in mouse models of MLLr leukemia (18). IVIS imaging studies were performed weekly and these documented near uniform engraftment and outgrowth of tumors in control mice. By Day 33 all control mice had evidence of disseminated disease (**Figure 7B-D**). By contrast, tumor formation was both delayed and highly variable in VTP50469-treated mice (**Figure 7B**). Half of the drug-exposed mice showed no evidence of tumor by IVIS imaging 40 days after cell injection (**Figure 7C**). This difference in tumor colonization between the groups led to significantly improved survival of drug-exposed mice (**Figure 7E**). On day 40, all but four treated mice were euthanized for necropsy. In the four surviving mice, drug was stopped and mice were followed for an additional 40 days, or humane endpoint whichever came first, and then necropsies performed.

**Figure 7.**
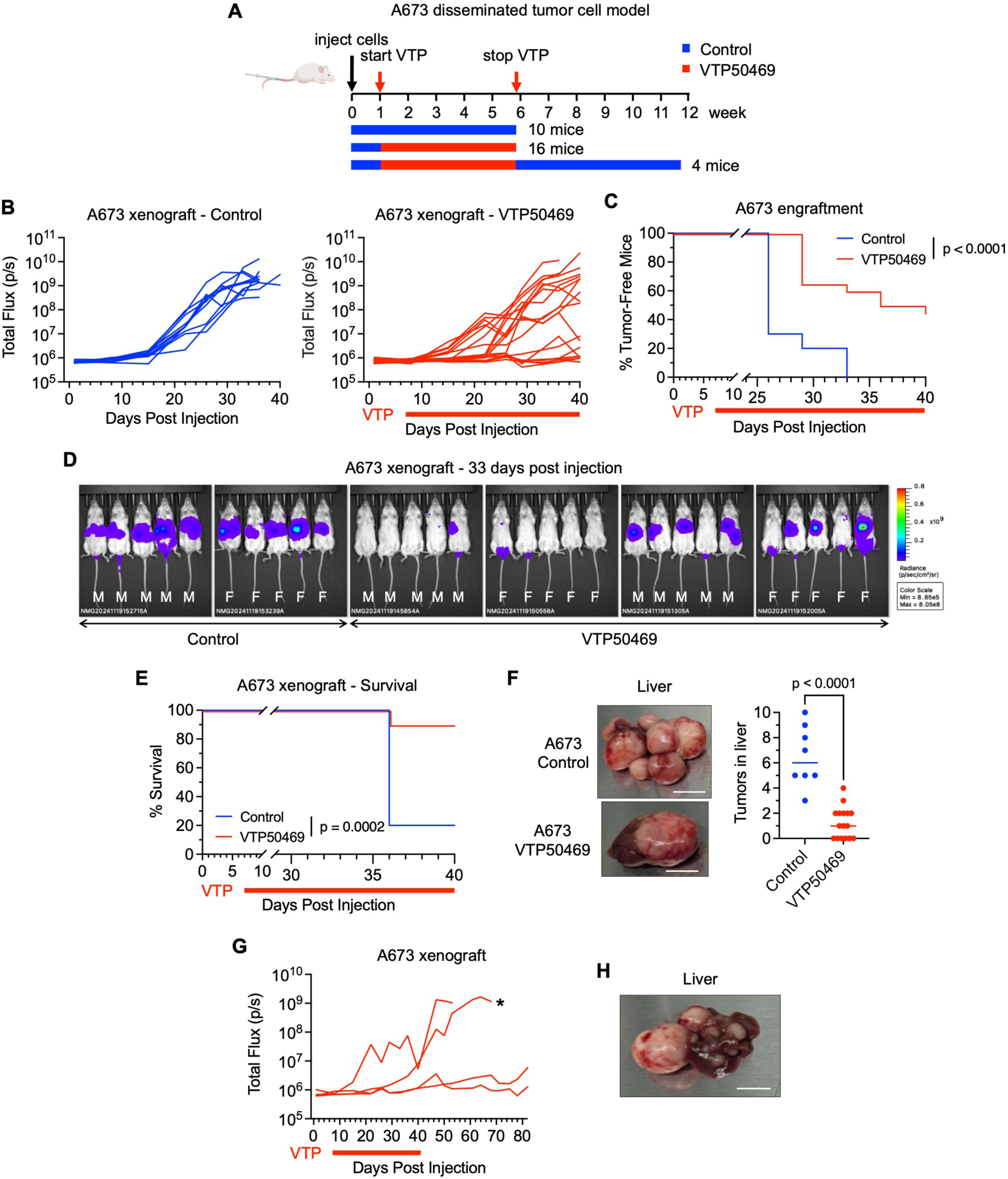
Menin inhibition reduces colonization in EwS xenografts. (**A**) Diagram of mouse experiment. 5e4 A673 GFP-Luciferase cells were injected into the tail veins of 30 NSG mice (male and female) and 7 days later mice randomized to regular (control) or 0.1% VTP50469 mouse chow (∼120-180 mg/kg/day). Four VTP50469-treated mice were taken off drug at study end and followed for up to 41 days as described in text. (**B**) Spider plots of IVIS signal for each control and VTP50469-treated mouse. (**C)** Plot of time to engraftment with engraftment set at an IVIS signal of 1e8. (**D**) Luminescence of all mice at 33 days post-injection. Sex of each mouse is indicated (M=male, F=female). (**E**) Survival plot comparing the control and VTP50469-treated mice. (**F**) Images of representative livers from the control and VTP50469-treated mice are shown (scale bar=1 cm). The number of macroscopic tumors in the livers was counted and plotted for each mouse. (**G**) Spider plot of IVIS signals from 4 mice taken off VTP50469 and followed. The liver of one of the released mice (* in **G**) is shown.

Necropsies of the mice euthanized on day 40 showed that tumors had mainly engrafted in the liver, consistent with prior studies of EwS in NSG models. Significantly, whereas multiple macroscopic tumors were clearly visible in the livers of all control mice, tumor burden was markedly reduced in VTP50469-treated mice, and in 6/16 mice no tumors could be identified (**Figure 7F**). Of the four mice that were observed after stopping drug, two failed to develop tumors after 40 days of drug-free follow-up (**Figure 7G**). Conversely, two mice developed liver masses and, in both mice, an increase in IVIS signal was evident at the time of drug cessation, presaging formation of macroscopic tumors (**Figure 7G**). Intriguingly, in one of the mice that developed tumors off drug, necropsy revealed that the liver contained a single very large tumor in addition to multiple much smaller tumors (**Figure 7H**). We speculate that these smaller tumors emerged only after VTP50469 was stopped, supporting a role for the Menin inhibitor in preventing outgrowth of micrometastatic foci.

To test if the inhibitory effects of VTP50469 on metastatic colonization would extend to an additional model, we repeated the experiment with TC32 cells (**Supplemental Figure 5A**). In this experiment, we followed half of the mice (N=10) for 25 days after drug removal to monitor tumor outgrowth after treatment cessation. As with A673 cells, metastatic colonization of disseminated TC32 cells was inhibited by VTP50469 and macroscopic tumor burden in livers after 6 weeks of drug treatment was significantly reduced (**Supplemental Figure 5B and C**). Intriguingly, the impact of the drug was much more significant in male compared to female mice largely because colonization of TC32 cells in control mice was more rapid and efficient in males than females (**Supplemental Figure 5C**). This was not observed with A673 cells (**Supplemental Figure 5D**) and the reason for this sex discrepancy in the TC32 model is unknown. Both A673 and TC32 cell lines were originally derived from female patients with localized tumors. Nevertheless, regardless of mouse sex, removal of VTP50469 led to rapid outgrowth of macroscopic tumors in all mice within 25 days and livers of both male and female mice were nearly completely replaced by tumor (**Supplemental Figure 5E**). Thus, pharmacologic inhibition of Menin using a Menin:MLL inhibitor reproducibly inhibited EwS tumor colonization. In most instances, withdrawing the drug resulted in rapid tumor outgrowth, supporting the hypothesis that inhibiting Menin activity reversibly suppresses the capacity of disseminated EwS cells to form tumors in metastatic niches.

## Discussion

Despite a robust and most often complete clinical response to primary therapy, patients with both localized and metastatic EwS often experience tumor relapse months or years after their original diagnosis (24). Metastatic relapse remains the primary cause of EwS patient death yet very little is understood about the mechanisms that underlie this enduring clinical problem and investigational agents have thus far offered no improvement in outcomes (33). Metastasis is a complex, multi-stage process that begins with invasion of cells beyond the confines of a primary tumor, continues with transit and survival of cells through the circulation, and culminates with engraftment and outgrowth of tumors at metastatic sites (34). Outgrowth of microscopic tumor foci into macroscopic metastases, or colonization, can take years and is the final and most inefficient step in the metastatic cascade (35). In the current work, we have identified the scaffolding protein Menin as a key driver of EwS metastasis.

Both genetic and pharmacologic loss of function approaches revealed that despite having little impact on cell phenotypes in vitro, inhibiting Menin activity in EwS cells significantly impairs their capacity to metastasize in vivo. Moreover, this was found to be primarily mediated by a defect in tumor colonization.

In both normal development and in cancer, Menin functions to activate and repress gene transcription via its highly context-specific interactions with diverse partner proteins, including transcription factors and chromatin modifying enzymes (2, 22). The most well-studied and ubiquitous interaction partners of Menin are the MLL1 and MLL2 histone methyltransferases, and our studies show that MLL2 robustly interacts with Menin in EwS cells. Disruption of Menin:MLL interactions can be achieved by pharmacologic inhibitors of Menin that were developed for treatment of MLLr leukemias (21). Here, we have shown that VTP50469 broadly alters gene expression profiles of EwS cells and this is associated with disruption of Menin:MLL2 interactions. Intriguingly, however, genes that were most reproducibly downregulated by genetic or pharmacologic inhibition of Menin in EwS cells were MYC target genes. In addition, we found that Menin forms a complex with MYC in EwS and that Menin:MYC interactions are partially disrupted by VTP50469. These data, combined with the more extensive changes to cell phenotype and gene expression that we identified in MEN1-KO compared to VTP50469-treated cells, lead us to conclude that Menin contributes to the pathogenesis of EwS via mechanisms that extend beyond MLL-dependent gene regulation. Significantly, Menin inhibitors have shown activity in preclinical models of other solid tumors (36–40) and clinical trials of these drugs in cancers beyond leukemia are now underway (NCT06655246 (gastrointestinal stromal tumors), NCT05731947 (colorectal cancer and solid tumors)). It remains to be elucidated whether the anti-tumor effects of Menin inhibitors in solid tumors are mediated in whole or in part by disrupting Menin interactions with MLL proteins or if direct or allosteric effects on other interaction partners, including MYC, are important. Given its broad and context-dependent partners and functions in normal organogenesis and development, we propose that the mechanisms by which Menin inhibitors impede oncogenic states in cancer will be, at least in part, tumor-type dependent.

In addition to downregulation of MYC signature genes, our studies revealed that genetic loss or pharmacologic inhibition of Menin led to upregulation of developmental, hypoxia, and stress-response programs. Notably, the specific genes and pathways that were suppressed by Menin expression and activity differed between A673 and TC32 cells, again demonstrating the highly cell context-dependent nature of Menin-regulated programs. Nevertheless, genes that control EMT were over-represented among Menin-repressed genes in both cell lines and in both genetic and pharmacologic loss of function studies. These findings in a sarcoma are consistent with the critical role for Menin in regulating development of mesoderm and neural crest tissues during organogenesis and mesenchymal lineage differentiation (3, 5–8). Importantly, over-expression of Menin suppresses terminal differentiation of osteoblasts and this is mediated by its antagonism of the JUND transcription factor (6). JUND interacts with Menin in the same binding pocket as MLL and this interaction blocks JNK/ERK2-mediated phosphorylation of JUND (41, 42). Given that Menin:MLL inhibitors also disrupt Menin:JUND interactions, we hypothesize that the biologic and transcriptomic effects of Menin loss of function on mesenchymal gene programs in EwS cells may be, in part, secondary to effects on JUND activity. In addition, EWS::ETS fusion proteins are the oncogenic drivers of EwS and are required for tumor initiation and maintenance (23). EWS::ETS proteins directly and indirectly repress expression of mesenchymal differentiation genes and loss of this repressive activity promotes migratory and metastatic cell phenotypes (29, 43–46). Thus, it is possible that Menin also promotes metastatic colonization of EwS cells by modifying the transcriptional activity of the EWS::ETS driver. Experiments to test this hypothesis are ongoing.

In summary, these studies identify Menin as a master regulator of gene transcription in EwS and a critical mediator of tumor metastasis. Our preclinical studies with the Menin inhibitor revumenib suggest that Menin:MLL inhibitors warrant investigation as novel agents for patients with high risk EwS. We propose that Menin inhibition could be leveraged to delay or prevent metastatic relapse in patients who are suspected to harbor subclinical residual disease at the end of primary therapy. Additional studies are now needed to determine if and how other agents could be combined with Menin inhibitors to eliminate rather than suppress growth of residual disease.

## Methods

### Sex as a biologic variable

A mix of male and female mice were used in the in vivo experiments described below to control for sex differences in tumor engraftment and metastasis. In VTP50469 studies, mice were randomized to treatment groups, stratifying to include both male and female mice. The sex of each mouse is indicated in the figures.

### Cell culture

A673 and TC32 EwS cell lines were grown in RPMI (Gibco 11875093) with 10% FBS (Atlas Biologicals EF-0500-A) and 1% L-glutamine (Gibco 25030081), CHLA10 EwS cell line was grown in IMDM (Gibco 12440053) with 20% FBS, 1% L-glutamine and 1X Insulin-Transferrin-Selenium-Ethanolamine (ITS-X, Gibco 51500056) and U2OS (osteosarcoma) cells were grown in McCoy’s 5A (Gibco 16600082) with 10% FBS, 1% L-glutamine at 37°C with 5% CO_2_. Cells were washed with Dulbecco’s phosphate-buffered saline (DPBS, Gibco 14040117) and passaged using 0.05% Trypsin-EDTA (Gibco 25300120). The CRISPR/Cas9 system was used to knock out Menin using guide 1 (gMEN1 1) to exon 2 (GCTGCGCTCCATCGACGACG) or guide 5 (gMEN1 5) to exon 3 (CAAATTGGACAGCTCCGGTG) and a nontargeting guide (gControl, GGTGGACATCCCGGAGACCC) for the control using the LentiCRISPRv2 vector (Addgene plasmid #52961). A673 and TC32 cells were transduced with each CRISPR lentivirus, transduced cells selected with puromycin, single cell clones isolated and verified for loss of Menin protein by western. The following CRISPR clones were generated with the last number in the name indicating the clone number (A673 gControl 1, A673 gMEN1 1-3, A673 gMEN1 5-1, gMEN1 5-2, TC32 gControl 3, TC32 gMEN1 1-5, TC32 gMEN1 5-5) and all were grown in the same conditions as the parental cell line. For animal experiments, cell lines were transduced with a GFP-Luciferase vector (pj01668-16revised-egfp-ffluc_ephiv7). To inhibit Menin, cells were treated with 0.1% DMSO (Vehicle) or 10 µM VTP50469 (MedChemExpress HY-114162) for 72 hours.

### RT-qPCR assay

Cells were trypsinized, pelleted, washed with ice-cold DPBS (Gibco 14040117) and total RNA (DNA-free) purified using the QIAshredder (Qiagen 79656), RNeasy Mini Kit (Qiagen 74106) and RNase-Free DNase Set (Qiagen 79254). cDNA was generated with iScript cDNA Synthesis Kit (Bio-Rad 1708891). qPCR reactions were done using iTaq Universal SYBR Green Supermix (Bio-Rad 1725122) with primers for *MYC* (TACAACACCCGAGCAAGGAC and GAGGCTGCTGGTTTTCCACT), *EEF1A1* (CGTTACAACGGAAGTAAAATC and CAGGATAATCACCTGAGC) and *18S* (GCAATTATTCCCCATGAACGA and GGCCTCACTAAACCATCCAAT) or TaqMan Fast Universal PCR Master Mix (Applied Biosystems 4352042) with primers for *MEN1* (Hs00365720_m1), *B2M* (Hs_00984230_m1) and *18S* (Hs_03003631_s1) and run on the Roche LightCycler 480 II. *MYC* Cp values were normalized to the geomean of the control primers Cp values (*EEF1A1* and *18S*) and *MEN1* Cp values were normalized to the geomean of the control primers Cp values (*B2M* and *18S*) and plotted as 2^ΔCp for Expression or relative to Cp value of the control cell sample normalized to the geomean of the control primers Cp values for Relative Expression (2^ΔΔCp).

### Western blot analysis

Cells were trypsinized, pelleted, washed with ice-cold DPBS (Gibco 14040117) and whole cell extracts were generated with a 10-minute incubation on ice in RIPA buffer (ThermoFisher 89900) with protease inhibitors (Roche 04 693 116 001) and phosphatase inhibitors (Roche 04 906 837 001), sonicated for 10×30 second cycles in the Bioruptor Pico (diagenode) and debris removed by centrifuging for 5 minutes at 16,000xg. Protein extracts were quantitated using the Bio-Rad Protein Assay Kit II (5000002) and 30 µg of total protein was loaded on 4–15% Mini-PROTEAN TGX Precast Protein Gels (Bio-Rad 4561084 or 4561086) with a Tris-Glycine-SDS buffer system (Bio-Rad 1610732) using Precision Plus Protein Dual Color Standards (Bio-Rad 161037) for molecular weight estimation. Proteins were transferred to a nitrocellulose membrane (Bio-Rad 1620115) using the Bio-Rad Mini Trans-Blot Electrophoretic Transfer Cell with Tris-Glycine buffer system (Bio-Rad 1610734).

Membranes were blocked with Intercept (TBS) Blocking Buffer (LICORbio 927-60001) and probed for the primary antibodies GAPDH Rabbit mAb (Cell Signaling 2118), MAX Rabbit pAb (Cell Signaling 4739), Menin Goat pAb (Bethyl A300-106A), MLL1-C (C-terminal) Rabbit pAb (Bethyl A300-374A), MLL2-C (C-terminal) Rabbit mAb (Cell Signaling 63735), MYC Rabbit mAb (Cell Signaling 18583), Vinculin Rabbit mAb (Cell Signaling 13901) or WDR5 Rabbit pAb (Bethyl A302-430A) followed by the secondary antibody Goat anti-Rabbit 800CW (Licor 926-32211), Donkey anti-Rabbit 800CW (Licor 926-32213) or Donkey anti-Goat 680RD (Licor 926-68074). All antibodies were diluted according to the manufacturer recommendations in Intercept (TBS) Blocking Buffer with 0.1% Tween-20. Membranes were scanned with the Odyssey (LICORbio) in the 700 and 800 nm channel.

### Mouse experiments

#### Subcutaneous mouse model

Cells were prepared by trypsinizing, washing with DPBS and resuspending in DPBS at 1e6 cells/50 µl, diluting 2-fold with 50 µl of Matrigel matrix (Corning 354234) and injecting 1e6 cells into the flank region on the right or left side just below the skin layer in a 7-week-old NSG mouse (NOD.Cg-Prkdcscid Il2rgtm1Wjl/SzJ) from Jackson Laboratory (strain # 005557).

Three to 5 mice were used for each cell line. Tumors were measured 2-5x per week with calipers in two dimensions, the volume calculated (V=(L x W2)/2) was plotted over time for each mouse. Once the first mouse reached endpoint, all mice were euthanized at the same time.

#### Tail vein mouse model

Cells transduced with the GFP-Luciferase vector were prepared by trypsinizing, washing with DPBS and resuspending in DPBS at 1e6 cells/100 µl and injecting 1e6 cells into the tail vein of a 7-week-old NSG mouse. Five mice were used for each cell line. Imaging with the In Vivo Imaging System (IVIS) was performed at 1 hour to confirm successful injection and then every 3-4 days until first mouse reached endpoint and then all mice were euthanized at the same time.

#### Subrenal capsule spontaneous metastasis mouse model

Cells transduced with the GFP-Luciferase vector were prepared by trypsinizing, washing with DPBS and resuspending in DPBS at 2e5 cells/5 µl, diluting 2-fold with 5 µl of Matrigel matrix. Live ultrasound imaging was used to insert the needle through the skin into the back muscle and into the subrenal capsule space to inject 2e5 cells into a 7-week-old NSG mouse (32). Five mice were used for each cell line. Tumor size in kidney was monitored by ultrasound and IVIS imaging was done 2x per week. Once the tumor in the kidney reached about half the size of the kidney in one mouse, all the mice were euthanized at once.

#### Disseminated tumor cell (DTC) mouse model

Cells transduced with the GFP-Luciferase vector were prepared by trypsinizing, washing with DPBS and resuspending in DPBS at 5e4 cells/100 µl and injecting 5e4 cells into the tail vein of 30 7-week-old NSG mice. After 7 days, 20 mice were switched to 0.1% VTP50469 mouse chow (∼120-180 mg/kg/day, SYNDAX PHARMACEUTICALS, INC.). Imaging with IVIS was done 1-2x per week. Treatment ended when last Control mouse reached endpoint (A673), or all control mice had a IVIS signal greater than 1e10 (TC32). The VTP50469-treated mice were either euthanized when endpoint was reached or at end of treatment with a total of 16 mice in A673 and 10 mice in TC32. Four (A673) or 10 (TC32) VTP50469-treated mice were switched to normal mouse chow to remove drug and monitored by IVIS imaging for ∼6 weeks (A673) or ∼3 weeks (TC32) at which point any remaining mice that did not reach endpoint were euthanized.

#### Tissue histology

Formalin-fixed, paraffin-embedded (FFPE) organs (liver and kidney) from the xenograft mouse experiments were sectioned, mounted on slides, deparaffinized and stained with hematoxylin and eosin (H&E). For immunofluorescence, following antigen retrieval, slides were blocked in blocking buffer (0.2% bovine serum albumin in PBS) for 1 hour, washed, incubated with 1:100 primary antibody, CD99 rabbit mAb (Biocare Medical CME392A), in blocking buffer for 1 hour, washed, incubated with 1:200 secondary antibody, Goat anti-Rabbit Alexa Fluor 647 (Invitrogen A-21245), in blocking buffer for 1 hour with 1:2000 DAPI (2.5 µg/ml, Invitrogen D1306) and coverslips were mounted with ProLong Gold Antifade Mountant (Invitrogen P36934). Whole slides were scanned with ZIESS Axioscan 7 or imaged with Leica DMi8 Thunder Imager at 20X magnification.

#### mRNA-sequencing

RNA was purified from cells of biological triplicates as described above and sent to Novogene for polyA enrichment library preparation using NEBNext Ultra II RNA Library Prep kit and sequencing with the Illumina Novaseq6000 sequencer utilizing a paired-end 150 bp (PE150) sequencing strategy. The quality of the sequencing data was analyzed with FastQC, adaptor fragments, low quality ends and short sequences were removed with Trim Galore, trimmed sequences were aligned to the reference genome, GRCh38, and gene counts generated with STAR, additional quality control was performed with MultiQC and differentially expressed genes were identified with DESeq2. Differentially regulated genes with a padj<0.05 were used to generate Venn diagrams using Vennerable. Hallmark and Gene Ontology Biological Processes (GOBP) pathway enrichment in commonly regulated genes was assessed using GSEA (47, 48), MSigDB (49, 50) or Enrichr (51). Heatmaps of differentially expressed genes were produced with pheatmap.

#### Real-time cell proliferation assays

Cells were plated in a 96-well plate with 5000 or 15,000 cells in 50 µl media per well and incubated overnight to adhere to plate. Media was added to 200 µl final and DMSO or VTP50469 were added to a final concentration of 0.1% and 10 µM, respectively, for the Menin inhibition assays. DPBS was added to the empty wells and interspaces in the plate to keep plate humidified. The plate was loaded into the Incucyte SX5 Live-Cell Analysis System (Sartorius) and images were captured every 2 hours. Confluence was measured using the IncuCyte software and plotted as a percentage over time.

#### 3D-collagen invasion assays

Small spheroids (10-1000 cells) were formed by trypsinizing cells, resuspending at 50,000 cells/ml, plating 4 ml/well in a non-adherent 6-well plate (Corning 3471) and incubating overnight. Collagen gels were formed by combining 1.8 mg/ml rat tail collagen I (Gibco A1048301) in 1X DMEM (Sigma-Aldrich D2429) and 8 mM acetic acid, neutralizing the mixture with 0.5 M NaOH and incubating on ice for 45-60 minutes to allow polymerization. A small flat circular underlay was formed by pipetting 25 µl of polymerized collagen mixture into each well of a 24-well plate warmed to 37°C. Spheroids embedded in collagen were formed by resuspending 1 ml of spheroids collected by centrifuging at 300xg for 10 seconds in 100 µl of polymerized collagen mixture, pipetting 60 µl of spheroid-collagen mixture on top of overlay, incubating at 37°C for ∼30 minutes to set gel and adding appropriate culture media. For VTP50469-treated cells, 0.1% DMSO or 10 µM VTP50469 were added with the media after the gels were formed. Plates were incubated for 5 days and then cells were fixed with 4% paraformaldehyde in PBS for 15 minutes, washed with PBS, permeabilized and stained with 0.1% Triton-X100, 1:2000 DAPI (2.5 µg/ml) and 1:800 Alexa Fluor 568 Phalloidin (400x, Invitrogen A12380) for 2 hours, washed with PBS and imaged in PBS on Leica DMi8 Thunder Imager at 10X magnification. Images were analyzed in Fiji by outlining spheroids and calculating circularity.

#### Co-immunoprecipitations

Menin (rabbit pAb, Bethyl Labs A300-105A), MYC (mouse mAb, Santa Cruz SC-42), Rabbit IgG, polyclonal - Isotype Control (Abcam ab37415) or Mouse IgG1, kappa monoclonal - Isotype Control (Abcam ab18443) were crosslinked to Protein A Dynabeads (Invitrogen 10001D) with 20 mM dimethyl pimelimidate dihydrochloride (Millipore Sigma D8388) at 8 µg human IgG/mg beads. Antibody-beads were resuspended at 30 mg/ml.

The nuclear extract was prepared using the Nuclear Complex Co-IP Kit (Active Motif 54001). Briefly, cells were trypsinized, washed in PBS with Phosphatase Inhibitors, resuspended in hypotonic buffer, lysed with Detergent, nuclei were pelleted by centrifuging at 10,000xg, nuclei were lysed in Digestion Buffer with Protease Inhibitor Cocktail and 0.5 mM PMSF and chromatin sheared with the Enzymatic Shearing Cocktail, enzymes were inactivated with 10 mM EDTA, debris removed by centrifuging at 12,000xg and the supernatant collected for the nuclear extract. Nuclear extracts were quantitated using the Bio-Rad Protein Assay Kit II.

For each immunoprecipitation, 1-2 mg of nuclear extract were incubated with 0.25 mg beads (2 µg Ab) per mg of nuclear extract in IP buffer (20 mM HEPES•KOH (pH 7.5), 150 mM NaCl, 0.5% Igepal, protease inhibitors (Roche 04 693 116 001) and phosphatase inhibitors (Roche 04 906 837 001)).

Immunoprecipitations were incubated on rotator at 4°C for 2 hours and washed 3X with IP Buffer and the protein was eluted with Laemmli Sample Buffer at room temperature for 10 minutes. Eluted samples and 5% and 1% of input (nuclear extract) prepared in Laemmli Sample Buffer were heated at 95°C for 5 minutes. Input and 50% of the eluted sample were analyzed by Western blots as described above.

#### Statistics

For cell culture experiments, graphical data is plotted as the mean with a standard deviation of at least 3 biological replicates and a Welsh’s t-test with a two-tailed p value at 95% confidence level was used to compare cell lines or treatment. For Venn diagrams, the GeneOverlap R package was used to calculate the p value of the overlap using a Fisher’s exact test. For animal experiments, time to engraftment and survival plots used a Logrank (Mantel-Cox) test to calculate the p value. The p values for the subcutaneous mouse experiments reported on the spaghetti plots are from the mean tumor size of mice at endpoint calculated with a Welsh’s t-test with a two-tailed p value at 95% confidence level.

The number of tumors in the DTC mouse experiments were plotted as the mean with standard deviation and a Welsh’s t-test with a two-tailed p value at 95% confidence level to compare treated to untreated mice. P values are displayed as follows: ns (nonspecific) p > 0.05, * p ≤ 0.05, ** p ≤ 0.01, *** p ≤ 0.001, **** ≤ 0.0001.

#### Study approval

All experimental procedures adhere to the *Guide for the Care and Use of Laboratory Animals* and were approved by Seattle Children’s Research Institute IACUC protocol ACUC00635. The institute is fully AAALAC accredited and has a Public Health Service approved Animal Welfare Assurance.

## Data availability

RNA-sequencing data will be submitted to NCBI Gene Expression Omnibus database upon publication,

## Author contributions

KAB and ERL conceived, designed, and supervised the study. KAB performed experiments, analyzed data and drafted the manuscript. NMG and DST performed animal experiments. MA, SIW, EDW and MEBD generated reagents, performed laboratory experiments, and analyzed data. ERL reviewed and analyzed data, edited the manuscript and secured funding for the study.

## Acknowledgments

The authors thank members of the Lawlor lab for helpful discussion, Gerard McGeehan from Syndax Pharmaceuticals, Inc. for supplying the VTP50469 mouse chow, Sean Taylor from the Seattle Children’s Research Institute Research Scientific Computing for assistance with the RNA-sequencing analysis and the staff at the Seattle Children’s Research Institute Microscopy and Histopathology CoLab and Office of Animal Care.

## Abbreviations

co-IP: co-immunoprecipitation experiments
DTC: disseminated tumor cells
ECM: Extracellular Matrix
EMT: Epithelial Mesenchymal Transition
EwS: Ewing sarcoma
IVIS: In Vivo Imaging System
MEN1-KO: *MEN1* knockout
MLLr: MLL rearranged
NPMm: NPM mutated
NSG: NOD scid gamma
VTP: VTP50469

## References

1. Matkar S, Thiel A, and Hua X. Menin: a scaffold protein that controls gene expression and cell signaling. Trends Biochem Sci. 2013;38(8):394–402.

2. Brown MR, and Soto-Feliciano YM. Menin: from molecular insights to clinical impact. Epigenomics. 2025;17(7):489–505.

3. Bertolino P, Radovanovic I, Casse H, Aguzzi A, Wang ZQ, and Zhang CX. Genetic ablation of the tumor suppressor menin causes lethality at mid-gestation with defects in multiple organs. Mech Dev. 2003;120(5):549–60.

4. Xu N, Cho HS, Hackland JOS, Benito-Kwiecinski S, Saurat N, Harschnitz O, et al. Genome-wide CRISPR screen identifies Menin and SUZ12 as regulators of human developmental timing. Nat Cell Biol. 2025;27(9):1411–21.

5. Engleka KA, Wu M, Zhang M, Antonucci NB, and Epstein JA. Menin is required in cranial neural crest for palatogenesis and perinatal viability. Dev Biol. 2007;311(2):524–37.

6. Naito J, Kaji H, Sowa H, Hendy GN, Sugimoto T, and Chihara K. Menin suppresses osteoblast differentiation by antagonizing the AP-1 factor, JunD. J Biol Chem. 2005;280(6):4785–91.

7. Sowa H, Kaji H, Canaff L, Hendy GN, Tsukamoto T, Yamaguchi T, et al. Inactivation of menin, the product of the multiple endocrine neoplasia type 1 gene, inhibits the commitment of multipotential mesenchymal stem cells into the osteoblast lineage. J Biol Chem. 2003;278(23):21058–69.

8. Sowa H, Kaji H, Hendy GN, Canaff L, Komori T, Sugimoto T, et al. Menin is required for bone morphogenetic protein 2- and transforming growth factor beta-regulated osteoblastic differentiation through interaction with Smads and Runx2. J Biol Chem. 2004;279(39):40267–75.

9. Inoue Y, Hendy GN, Canaff L, Seino S, and Kaji H. Menin interacts with beta-catenin in osteoblast differentiation. Horm Metab Res. 2011;43(3):183–7.

10. Yokoyama A, Somervaille TC, Smith KS, Rozenblatt-Rosen O, Meyerson M, and Cleary ML. The menin tumor suppressor protein is an essential oncogenic cofactor for MLL-associated leukemogenesis. Cell. 2005;123(2):207–18.

11. Hughes CM, Rozenblatt-Rosen O, Milne TA, Copeland TD, Levine SS, Lee JC, et al. Menin associates with a trithorax family histone methyltransferase complex and with the hoxc8 locus. Mol Cell. 2004;13(4):587–97.

12. Larsson C, Skogseid B, Oberg K, Nakamura Y, and Nordenskjold M. Multiple endocrine neoplasia type 1 gene maps to chromosome 11 and is lost in insulinoma. Nature. 1988;332(6159):85–7.

13. Chandrasekharappa SC, Guru SC, Manickam P, Olufemi SE, Collins FS, Emmert-Buck MR, et al. Positional cloning of the gene for multiple endocrine neoplasia-type 1. Science. 1997;276(5311):404–7.

14. Lemmens I, Van de Ven WJ, Kas K, Zhang CX, Giraud S, Wautot V, et al. Identification of the multiple endocrine neoplasia type 1 (MEN1) gene. The European Consortium on MEN1. Hum Mol Genet. 1997;6(7):1177–83.

15. Marx SJ. Molecular genetics of multiple endocrine neoplasia types 1 and 2. Nat Rev Cancer. 2005;5(5):367–75.

16. Chen YX, Yan J, Keeshan K, Tubbs AT, Wang H, Silva A, et al. The tumor suppressor menin regulates hematopoiesis and myeloid transformation by influencing Hox gene expression. Proc Natl Acad Sci U S A. 2006;103(4):1018–23.

17. Grembecka J, He S, Shi A, Purohit T, Muntean AG, Sorenson RJ, et al. Menin-MLL inhibitors reverse oncogenic activity of MLL fusion proteins in leukemia. Nat Chem Biol. 2012;8(3):277–84.

18. Krivtsov AV, Evans K, Gadrey JY, Eschle BK, Hatton C, Uckelmann HJ, et al. A Menin-MLL Inhibitor Induces Specific Chromatin Changes and Eradicates Disease in Models of MLL-Rearranged Leukemia. Cancer Cell. 2019;36(6):660–73 e11.

19. Uckelmann HJ, Kim SM, Wong EM, Hatton C, Giovinazzo H, Gadrey JY, et al. Therapeutic targeting of preleukemia cells in a mouse model of NPM1 mutant acute myeloid leukemia. Science. 2020;367(6477):586–90.

20. Issa GC, Aldoss I, DiPersio J, Cuglievan B, Stone R, Arellano M, et al. The menin inhibitor revumenib in KMT2A-rearranged or NPM1-mutant leukaemia. Nature. 2023;615(7954):920–4.

21. Nadiminti KVG, Sahasrabudhe KD, and Liu H. Menin inhibitors for the treatment of acute myeloid leukemia: challenges and opportunities ahead. J Hematol Oncol. 2024;17(1):113.

22. Majer AD, Hua X, and Katona BW. Menin in Cancer. Genes (Basel). 2024;15(9).

23. Riggi N, Suva ML, and Stamenkovic I. Ewing’s Sarcoma. N Engl J Med. 2021;384(2):154–64.

24. Zollner SK, Amatruda JF, Bauer S, Collaud S, de Alava E, DuBois SG, et al. Ewing Sarcoma-Diagnosis, Treatment, Clinical Challenges and Future Perspectives. J Clin Med. 2021;10(8).

25. Jimenez JA, Apfelbaum AA, Hawkins AG, Svoboda LK, Kumar A, Ruiz RO, et al. EWS-FLI1 and Menin Converge to Regulate ATF4 Activity in Ewing Sarcoma. Mol Cancer Res. 2021;19(7):1182–95.

26. Svoboda LK, Bailey N, Van Noord RA, Krook MA, Harris A, Cramer C, et al. Tumorigenicity of Ewing sarcoma is critically dependent on the trithorax proteins MLL1 and menin. Oncotarget. 2017;8(1):458–71.

27. Svoboda LK, Teh SSK, Sud S, Kerk S, Zebolsky A, Treichel S, et al. Menin regulates the serine biosynthetic pathway in Ewing sarcoma. J Pathol. 2018;245(3):324–36.

28. Wu G, Yuan M, Shen S, Ma X, Fang J, Zhu L, et al. Menin enhances c-Myc-mediated transcription to promote cancer progression. Nat Commun. 2017;8:15278.

29. Franzetti GA, Laud-Duval K, van der Ent W, Brisac A, Irondelle M, Aubert S, et al. Cell-to-cell heterogeneity of EWSR1-FLI1 activity determines proliferation/migration choices in Ewing sarcoma cells. Oncogene. 2017;36(25):3505–14.

30. Apfelbaum AA, Wu F, Hawkins AG, Magnuson B, Jimenez JA, Taylor SD, et al. EWS-FLI1 and HOXD13 control tumor cell plasticity in Ewing sarcoma. Clin Cancer Res. 2022.

31. Wrenn ED, Apfelbaum AA, Rudzinski ER, Deng X, Jiang W, Sud S, et al. Cancer-Associated Fibroblast-Like Tumor Cells Remodel the Ewing Sarcoma Tumor Microenvironment. Clin Cancer Res. 2023;29(24):5140–54.

32. Thomas TT, Chukkapalli S, Van Noord RA, Krook M, Hoenerhoff MJ, Dillman JR, et al. Utilization of Ultrasound Guided Tissue-directed Cellular Implantation for the Establishment of Biologically Relevant Metastatic Tumor Xenografts. J Vis Exp. 2018(135).

33. Collier AB, 3rd, Krailo MD, Dang HM, DuBois SG, Hawkins DS, Bernstein ML, et al. Outcome of patients with relapsed or progressive Ewing sarcoma enrolled on cooperative group phase 2 clinical trials: A report from the Children’s Oncology Group. Pediatr Blood Cancer. 2021;68(12):e29333.

34. Gerstberger S, Jiang Q, and Ganesh K. Metastasis. Cell. 2023;186(8):1564–79.

35. Massague J, and Obenauf AC. Metastatic colonization by circulating tumour cells. Nature. 2016;529(7586):298–306.

36. Nair PR, Danilova L, Gomez-de-Mariscal E, Kim D, Fan R, Munoz-Barrutia A, et al. MLL1 regulates cytokine-driven cell migration and metastasis. Sci Adv. 2024;10(11):eadk0785.

37. Olsen SN, Anderson B, Hatton C, Chu Z, Simpkins C, Wen Y, et al. Combined inhibition of KAT6A/B and Menin reverses estrogen receptor-driven gene expression programs in breast cancer. Cell Rep Med. 2025;6(7):102192.

38. Malik R, Khan AP, Asangani IA, Cieslik M, Prensner JR, Wang X, et al. Targeting the MLL complex in castration-resistant prostate cancer. Nat Med. 2015;21(4):344–52.

39. Cherif C, Nguyen DT, Paris C, Le TK, Sefiane T, Carbuccia N, et al. Menin inhibition suppresses castration-resistant prostate cancer and enhances chemosensitivity. Oncogene. 2022;41(1):125–37.

40. Hemming ML, Benson MR, Loycano MA, Anderson JA, Andersen JL, Taddei ML, et al. MOZ and Menin-MLL Complexes Are Complementary Regulators of Chromatin Association and Transcriptional Output in Gastrointestinal Stromal Tumor. Cancer Discov. 2022;12(7):1804–23.

41. Huang J, Gurung B, Wan B, Matkar S, Veniaminova NA, Wan K, et al. The same pocket in menin binds both MLL and JUND but has opposite effects on transcription. Nature. 2012;482(7386):542–6.

42. Agarwal SK, Guru SC, Heppner C, Erdos MR, Collins RM, Park SY, et al. Menin interacts with the AP1 transcription factor JunD and represses JunD-activated transcription. Cell. 1999;96(1):143–52.

43. Chaturvedi A, Hoffman LM, Welm AL, Lessnick SL, and Beckerle MC. The EWS/FLI Oncogene Drives Changes in Cellular Morphology, Adhesion, and Migration in Ewing Sarcoma. Genes Cancer. 2012;3(2):102–16.

44. Pedersen EA, Menon R, Bailey KM, Thomas DG, Van Noord RA, Tran J, et al. Activation of Wnt/beta-Catenin in Ewing Sarcoma Cells Antagonizes EWS/ETS Function and Promotes Phenotypic Transition to More Metastatic Cell States. Cancer Res. 2016;76(17):5040–53.

45. Apfelbaum AA, Wrenn ED, and Lawlor ER. The importance of fusion protein activity in Ewing sarcoma and the cell intrinsic and extrinsic factors that regulate it: A review. Front Oncol. 2022;12:1044707.

46. Suresh V, Hafemeister C, Konstantinou A, Grissenberger S, Sturtzel C, Cidre-Aranaz F, et al. Dynamic modelling of EWS::FLI1 fluctuations reveals molecular determinants of phenotypic tumor plasticity and prognosis in Ewing sarcoma. bioRxiv. 2025:2025.04.03.647002.

47. Mootha VK, Lindgren CM, Eriksson KF, Subramanian A, Sihag S, Lehar J, et al. PGC-1alpha-responsive genes involved in oxidative phosphorylation are coordinately downregulated in human diabetes. Nat Genet. 2003;34(3):267–73.

48. Subramanian A, Tamayo P, Mootha VK, Mukherjee S, Ebert BL, Gillette MA, et al. Gene set enrichment analysis: a knowledge-based approach for interpreting genome-wide expression profiles. Proc Natl Acad Sci U S A. 2005;102(43):15545–50.

49. Liberzon A, Subramanian A, Pinchback R, Thorvaldsdottir H, Tamayo P, and Mesirov JP. Molecular signatures database (MSigDB) 3.0. Bioinformatics. 2011;27(12):1739–40.

50. Liberzon A, Birger C, Thorvaldsdottir H, Ghandi M, Mesirov JP, and Tamayo P. The Molecular Signatures Database (MSigDB) hallmark gene set collection. Cell Syst. 2015;1(6):417–25.

51. Xie Z, Bailey A, Kuleshov MV, Clarke DJB, Evangelista JE, Jenkins SL, et al. Gene Set Knowledge Discovery with Enrichr. Curr Protoc. 2021;1(3):e90.

