## Supplemental Figures for "Menin inhibition impairs metastatic colonization of Ewing sarcoma"

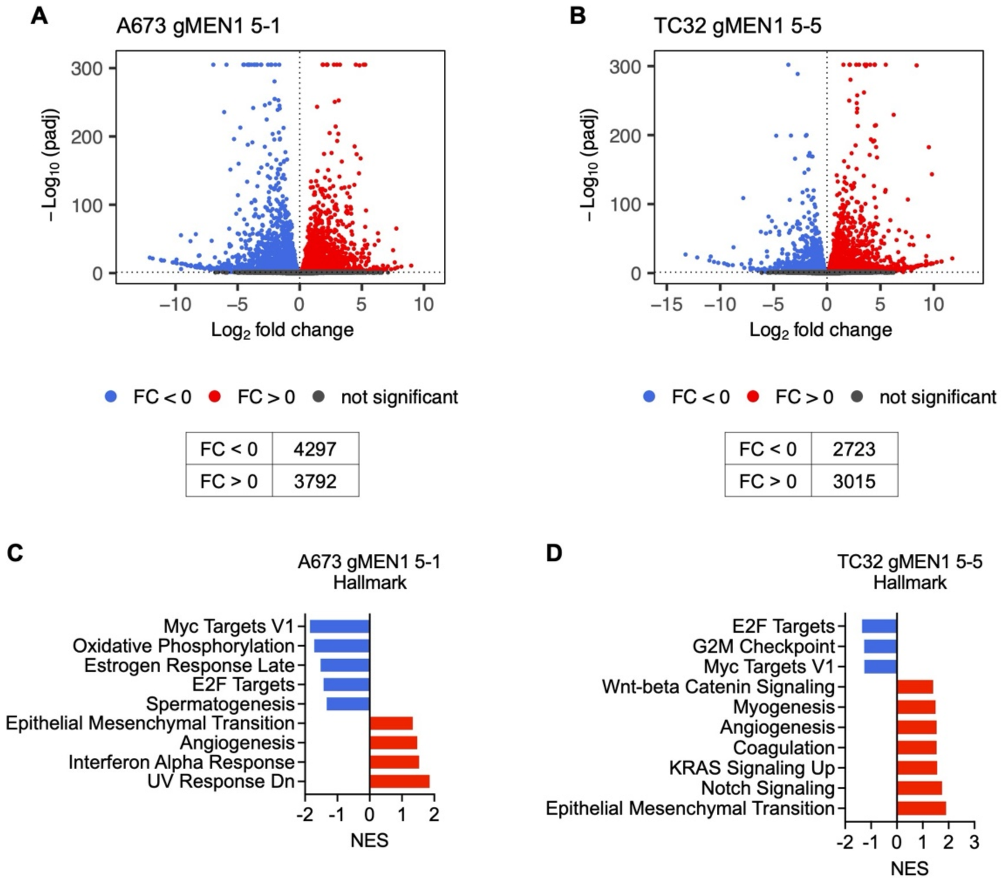


**Supplemental Figure 1**. **Menin regulates the transcriptome in A673 and TC32 cells.** (**A** and **B**) Volcano plots of differentially expressed genes between control and MEN1-KO clones using a significance cutoff of padj<0.05. The number of significantly downregulated (blue) and upregulated (red) transcripts are indicated in the tables. (**C** and **D**) Plots show most enriched Hallmark pathways among differentially expressed genes for each cell line (FDR q-val < 0.25).

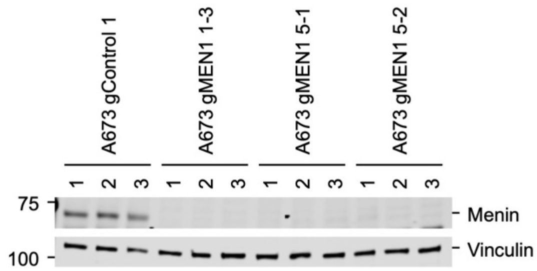


**Supplemental Figure 2. Loss of Menin protein is maintained in MEN1-KO tumors.** Western blots of total protein lysates isolated from subcutaneous xenografts show persistent loss of menin expression in all MEN1-KO cell-derived tumors.

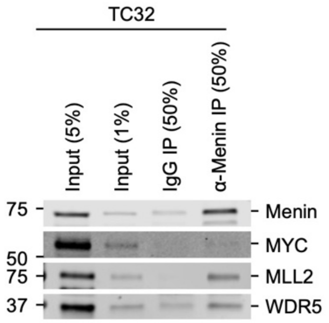


**Supplemental Figure 3. Menin and MYC co-immunoprecipitation in TC32.** Co-immunoprecipitations with a Menin antibody were performed on TC32 nuclear extract and immunoblotted for Menin, MYC, MLL2 and WDR5.

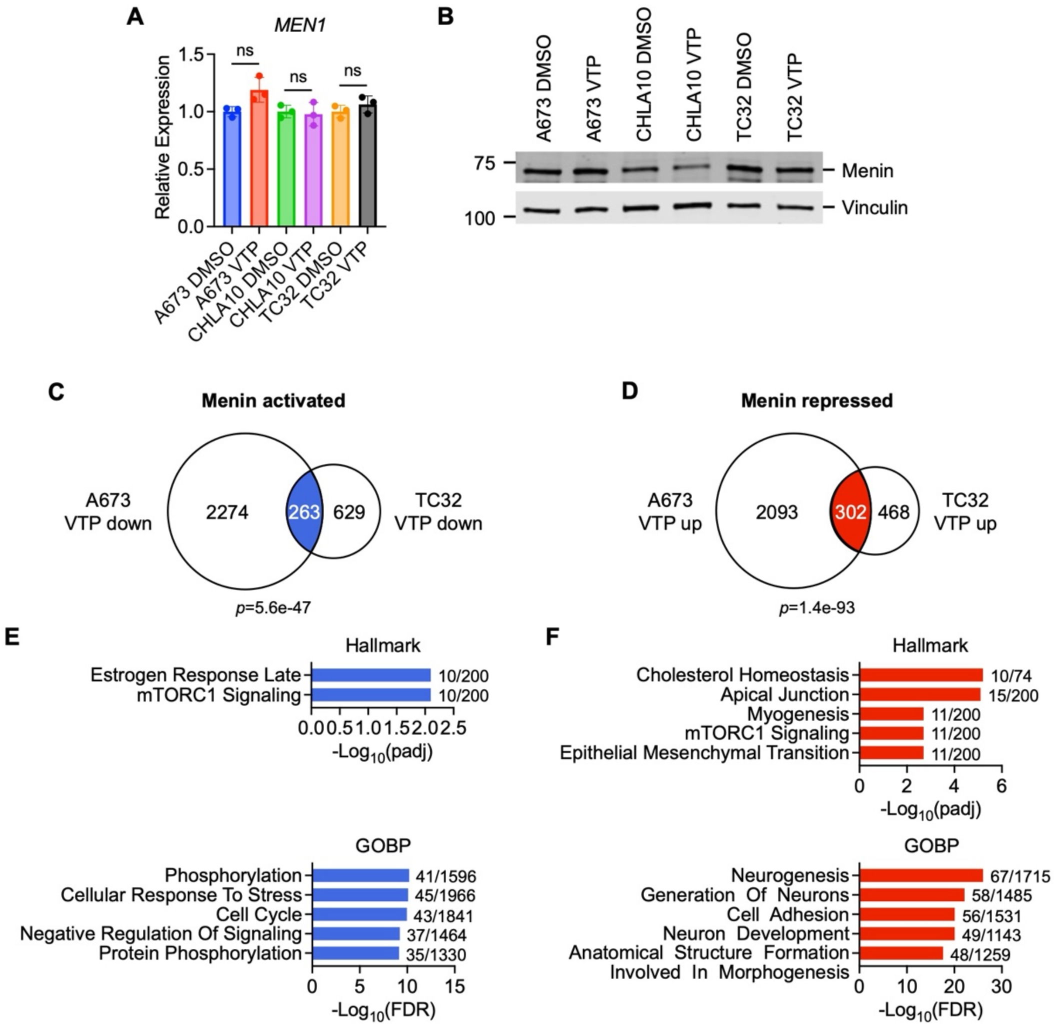


**Supplemental Figure 4. Menin inhibition regulates transcription independent of Menin levels.** (**A**) *MEN1* transcript levels in A673, CHLA10 and TC32 cells treated with 0.1% DMSO or 10 μM VTP50469 for 72 hours as determined by RT-qPCR. P values for VTP50469 vs DMSO were not significant for any of the cell lines (ns p > 0.05). (**B**) Western of Menin and Vinculin in A673, CHLA10 and TC32 cells treated with 0.1% DMSO or 10 μM VTP50469 for 72 hours. (**C** and **D**) A673 and TC32 cells were treated with 0.1% DMSO or 10 μM VTP50469 for 72 hours and genes that were significantly (padj<0.05) down-regulated (**C**) or up-regulated (**D**) in VTP50469-treated cells determined by RNA-seq. Overlap between the two cell lines is shown. (**E** and **F**) The enriched down-regulated (**E**) and up regulated (**F**) Hallmark pathways (Enrichr, padj<0.05) and top 5 GOBP enriched processes (MSigDB) are plotted with the overlap/total genes shown for each pathway.

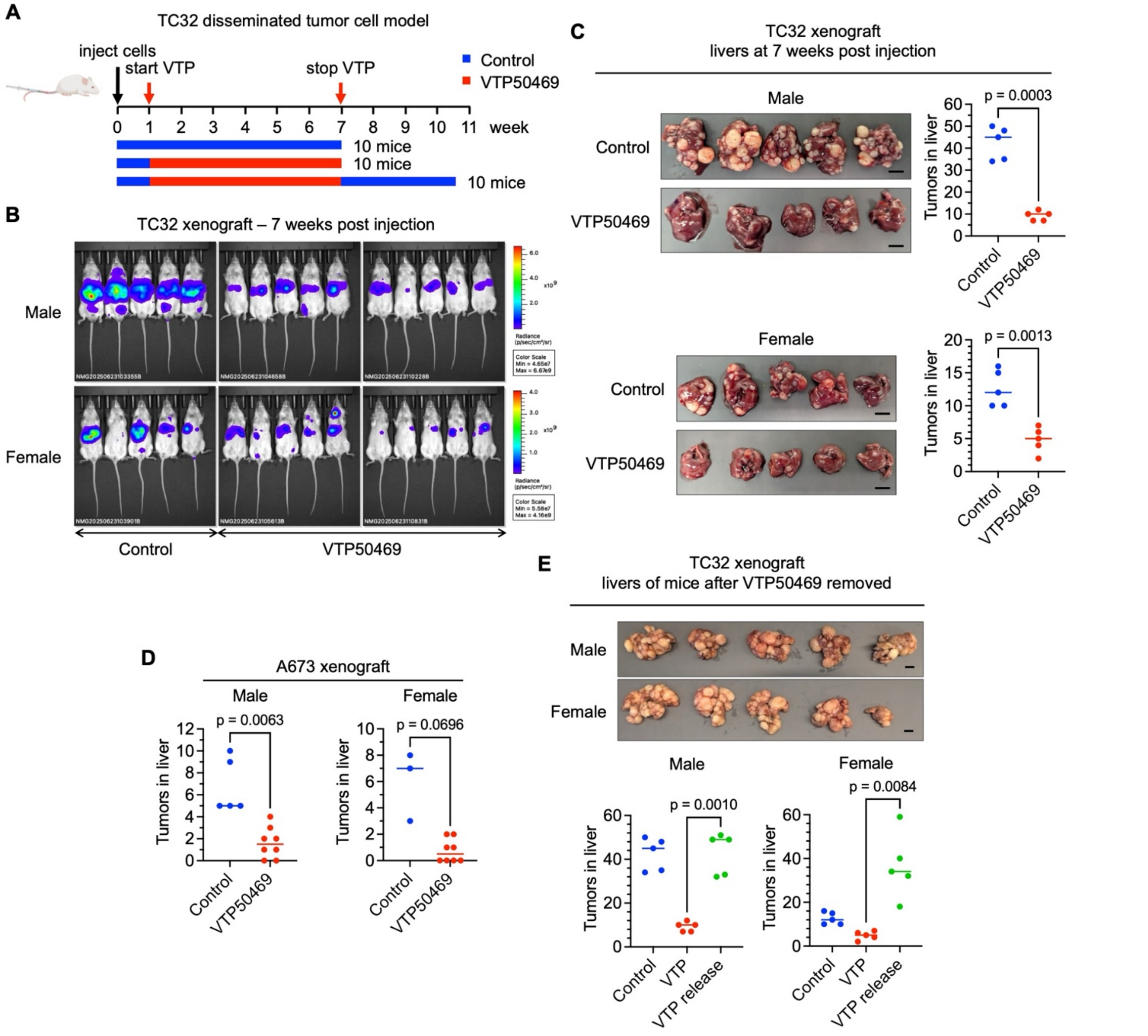


**Supplemental Figure 5. Menin inhibition reduces colonization in EwS xenografts.** (**A**) Diagram of mouse experiment. 5e4 TC32 GFP-Luciferase cells were injected into the tail vein of 30 NSG mice (male and female) and 7 days later 20 mice (VTP50469) were fed 0.1% VTP50469 mouse chow (~120-180 mg/kg/day) and 10 mice (control) were fed regular mouse chow. Seven weeks post injection, 10 VTP50469-treated mice were taken off drug and followed for up to 25 days. (**B**) IVIS images of the mice at 7 weeks post injection. (**C**) Livers of 10 control and 10 VTP50469-treated mice at necropsy 7 weeks post-cell inoculation. The number of macroscopic tumors in the livers were counted and plotted for each mouse. Males and females are shown separately to demonstrate sex differences. (**D**) Numbers of macroscopic tumors in male and female mice inoculated with A673 cells and then fed control or VTP50469 chow (from **Figure 7**). (**E**) Photos of livers taken at necropsy from 10 mice that were followed after removing VTP50469. The number of macroscopic tumors in the livers were counted and plotted for each mouse. Data from control and VTP50469-treated tumors (from **C**) are included for comparison.
